# From Home to Transcriptome: Comparing the transcriptomic profile of induced immune response via lipopolysaccharide stimulation in homeRNA and venous blood

**DOI:** 10.1101/2025.09.27.678977

**Authors:** Lauren G. Brown, Xiaofu Wei, Laura A. Milton, M. Yunos Alizai, James W. MacDonald, Theo K. Bammler, Yuting Zeng, Ingrid H. Robertson, Karen N. Adams, Yi-Chin Toh, Damien Chaussabel, Erwin Berthier, Amanda J. Haack, Ashleigh B. Theberge

## Abstract

Remote blood sampling offers multiple advantages over traditional clinic-based blood sampling studies, including greater patient inclusion, more frequent sampling, and broader geographical reach. Combining remote blood sampling with transcriptomic analysis opens potential in translational applications for capturing acute and dynamic immune responses to various exposures. In this study, we establish the feasibility of homeRNA, a capillary blood collection and RNA*later*-based stabilization kit, for use in downstream bulk RNA-sequencing applications via capturing a lipopolysaccharide (LPS)-induced inflammatory response. We also compared the baseline gene expression profiles and induced inflammatory response following LPS stimulation between homeRNA-stabilized samples and venous blood stabilized with RNA*later* or PAXgene. We found that homeRNA was successfully able to capture an inflammatory response to LPS, specifically targeting various cytokines (e.g., *IL6, IL12B, IL1B*), chemokines (e.g., *CCL3, CXCL10, CCL4*), and other transcriptional factors in the toll-like receptor pathway, the primary pathway activated during LPS stimulation. Importantly, we also found that homeRNA captured a LPS-induced inflammatory response comparable to that of venous blood samples stabilized with either RNA*later* or PAXgene. Overall, this work demonstrates that the homeRNA platform is compatible with downstream bulk RNA-sequencing analysis and can capture transcriptomic immune responses to a known stimulus which are analogous to results in traditional stabilized venous blood samples.

## INTRODUCTION

The convergence of remote blood sampling technologies and transcriptomic analyses enables large-scale, longitudinal gene expression studies that capture transient immune responses outside of the constraints of clinical settings. By eliminating the need for centralized clinic locations and trained phlebotomy staff, remote blood sampling not only increases geographical sampling locations, but also facilitates greater inclusion of participants who would otherwise be excluded due to logistical or personal constraints^1–5^. Further, with a remote study design, researchers can track immune responses to real-world exposures and collect frequent longitudinal samples without the burden of repeated clinic visits^6,7^.

Among the various molecular readouts possible from blood samples, RNA expression profiling is of particular value for capturing dynamic immune responses^8–10^. However, the inherent instability of whole blood RNA presents a significant challenge for *ex vivo* transcriptomic profiling; endogenous RNases can activate degradation pathways that compromise the integrity of intracellular transcripts, which can alter gene expression levels^11,12^. Therefore, studies that do not include immediate extraction of RNA from collected blood samples require a method of RNA stabilization. In clinic-based studies, commercially available RNA stabilization solutions such as PAXgene® and Tempus™ are commonly used; these solutions can come packaged in vacutainer tubes which make it easy to directly collect blood into the stabilizer solution from a phlebotomy draw. RNA*later*™ is another available stabilization solution which has been used to stabilize RNA in both blood and tissue samples^13,14^.

The need for RNA stabilization becomes even more critical when samples must be collected remotely and shipped to centralized laboratories. A commonly used method for remote sampling is dried blood spot (DBS)-based sampling, which typically uses a lancet on a fingertip to collect a drop of blood on a piece of paper^15,16^. DBS sampling relies on the drying of the blood sample to stabilize RNA. Many studies have successfully used RNA-sequencing on DBS samples^17–19^, but there is no standardized protocol for processing RNA from these samples^20–22^ and a pre-amplification step is often required prior to sequencing due to low RNA yield^17,23,24^. To increase RNA yield from remotely collected blood, there have been additional blood collection devices that utilize a lancet that collects blood from the upper arm (e.g., Tasso, YourBio) ^25,26^. These collection methods will typically draw up to 1 mL of blood, which is incompatible with drying, so liquid stabilization is required to preserve RNA integrity in transit.

To address this need, we developed the homeRNA kit, an at-home blood self-collection and RNA stabilization kit that uses the commercially available Tasso-SST device, an upper-arm blood collection device, and a custom stabilizer tube containing RNA*later*^27^. Blood samples collected with homeRNA have demonstrated sufficient RNA integrity number (RIN) value cutoffs for typical requirements for RNA sequencing analysis, suggesting that homeRNA allows for sufficient stabilization during the remote self-sampling, user-stabilization, and shipping processes^27^. Further, we demonstrated that homeRNA was robust during the shipping process (i.e., shipping time and exposure to high temperatures), in which we found negligible effects on subsequent gene expression using 3’ mRNA-sequencing^28,29^. Critically, we also demonstrated that homeRNA can effectively capture a biological immune response to SARS-CoV-2 using a targeted Nanostring gene panel^6,30^.

In this work, we demonstrate the applicability of homeRNA to detect a biological response with bulk RNA-sequencing transcriptomic analysis to build on our prior work with 3’ mRNA-sequencing^29^ and Nanostring analysis^6,30^. Specifically, we are interested in leveraging the larger read depth of bulk RNA-sequencing technologies in comparison to that of 3’ mRNA-sequencing and Nanostring analysis to investigate gene expression signatures in homeRNA-collected samples and standard phlebotomy blood draws. In this study, we measure an induced immune response to lipopolysaccharide (LPS)—a well-established immune stimulation model^31–33^—in homeRNA-collected and stabilized blood compared to RNA*later*-stabilized or PAXgene-stabilized venous blood. By comparing the gene expression profiles between homeRNA-stabilized capillary blood and traditional phlebotomy-drawn venous blood, this study establishes the compatibility of homeRNA with bulk RNA-sequencing technologies and demonstrates that homeRNA can capture a LPS-induced inflammatory response similar to that of venous blood. This study aims to provide critical guidance for researchers implementing homeRNA-based remote blood collection in studies investigating immune activation.

## METHODS AND MATERIALS

### Participant Recruitment and Demographics

Healthy adult volunteers (18 years or older) were recruited in Seattle, Washington via word of mouth under a protocol approved by the University of Washington Institutional Review Board study number STUDY00007868. Written informed consent was obtained from all participants (n=3). Each participant had 10 mL of venous blood collected into an EDTA-coated BD vacutainer (Becton Dickenson) as well as 0.6 mL to 1.5 mL of blood collected from the upper arm using two to three Tasso-SST devices, which was pooled together. The authors note that the serum separator tube (SST) gel (included in the Tasso-SST collection tube) is not necessary for RNA stabilization and analysis. At the time of the study, the Tasso-SST was the only available Tasso device for purchase.

### LPS stimulation and blood stabilization

Lipopolysaccharide from Escherichia coli O111:B4 (Sigma Aldrich) was reconstituted in RPMI culture media (Gibco) to make a 1 mg/mL stock solution. After blood collection, the blood collected from the two to three separate Tasso-SST™ devices used by a single participant was pooled into a microcentrifuge tube and mixed with a pipette until homogeneous. For each donor, 150 μL of venous blood and Tasso-collected blood was pipetted into separate replicates (n=2 replicates for each condition for a total of four samples for Tasso-collected blood and eight samples for venous blood per donor) within a 24-well plate. 150 μL of RPMI media containing LPS was added to each LPS stimulation replicate to achieve a final concentration of 100 ng/mL LPS. 150 μL of RPMI media (not containing any LPS) was added to each -LPS control replicate. Each sample was then incubated at 37°C for 6 h. Immediately after incubation, each venous blood sample (300 μL total including blood and added media volume) was transferred to microcentrifuge tube and stabilized according to manufacturer’s protocol with either RNA*later*™ (1:2.6 mL blood:RNA*later*™) or PAXgene® (1:2.76 mL blood:PAXgene®). For the Tasso-collected samples, each blood sample (300 μL total including blood and added media volume) was transferred back into a fresh Tasso-SST blood collection tube which was then attached to a custom engineered stabilizer tube used in the homeRNA kit, which contained 1.3 mL RNA*later*, and shaken to mimic blood stabilization in homeRNA kits^27^. After stabilization, blood samples were stored at -80°C until ready for RNA isolation.

### RNA isolation

For all blood samples stabilized with RNA*later*, total cellular RNA was isolated using the Ribopure Blood RNA Isolation Kit (Thermo Fisher) according to the manufacturer’s protocol without DNase treatment. Isolated RNA was stored at -80°C until ready for sequencing. For all blood samples stabilized with PAXgene, total cellular RNA was isolated using the PAXgene Blood RNA Kit (PreAnalytix) according to the manufacturer’s protocol without DNase treatment. Due to procedural errors, one donor’s PAXgene-stabilized samples were not usable.

### RNA sequencing and analysis

#### Library preparation and sequencing

Total RNA samples were shipped to Psomagen for library preparation and sequencing. Prior to library preparation, all samples were treated with DNase I according to manufacturer’s protocol. After DNase treatment, RNA quantity was measured using fluorescence-based quantification method Ribogreen (Life technologies) and RNA quality was assessed using Agilent RNA screentape (Agilent Technologies) on Agilent 4200 TapeStation system. All samples yielded RNA Quality Numbers (RQNs) ≥ 6.5 (see Supplementary Figure S1 for RQN distributions for each blood collection and stabilization condition). For each sample, 25ng of RNA was used as input for library preparation using Illumina Stranded Total RNA Prep with Ribo-Zero Plus kit. The resulting libraries were validated with D1000 Screen Tape (Agilent Technologies) and D1000 Reagents (Agilent Technologies) on 4200 TapeStation system (Agilent Technologies) to determine library size. Library quantification was measured with the Roche library quantification kit on LightCycler480 (Roche). Validated libraries were then normalized, diluted to the desired loading concentration and sequenced on the NovaSeq X Plus system using paired-end 151bp reads, targeting 15Gb per sample. Sequencing was performed using the NovaSeq X Plus system software. Real Time Analysis software was used to perform base calling from the raw images files generated by the Illumina sequencer. The resulting binary BCL/cBCL files were then converted to FASTQ files format using bcl2fastq, an Illumina provided package.

#### RNA sequencing quality control, alignment, and quantification

Quality of raw sequencing data was assessed using fastqc (https://www.bioinformatics.babraham.ac.uk/projects/fastqc/), and then reads were aligned to the NBCI GRCh38 transcriptome using the salmon aligner^34^. Supplementary Tables S1 and S2 detail the total reads obtained, number of reads aligned, and alignment percentage for each individual sample and group, respectively. Transcript abundances were read into R and summarized at the gene level using the Bioconductor tximport package^35^. Genes with unreliably low expression were excluded, with 19,751 genes retained for analysis.

#### Identification of differentially expressed genes

Differentially expressed genes were identified using the limma-voom pipeline^36^. Briefly, gene counts were adjusted for library size by computing counts per million counts (CPM), and then taking logs (logCPM). The logCPM values have reduced right skew, but still have a dependence between the mean and variance, which violates a core assumption for linear regression. Observation-level weights were estimated from the trend between the mean and variance and used in a weighted linear regression to account for the heteroscedasticity. Between-group comparisons were made using empirical Bayes adjusted contrasts, selecting genes at a false discovery rate (FDR) < 0.05, and incorporating an additional 30% change criterion in our statistic^37^.

#### Heatmap visualization

Heatmaps were generated in R (v4.5.1) using the pheatmap package (v1.0.13). Genes included in the heatmaps were selected as differentially expressed (FDR < 0.05) with at least 30% change in expression. Batch effects from donors were removed using the *removeBatchEffect* function in the limma package (v3.64.1)^38^. Z-scores, used for row scaling, were calculated from log₂-transformed counts per million (logCPM) values generated using the edgeR (v4.6.2) package.

#### Volcano plot visualization

Volcano plots were generated using iPathwayGuide™ (AdvaitaBio Corporation, https://ipathwayguide.advaitabio.com/)^39,40^.

#### Gene ontology enrichment and pathway analysis

Gene Ontology (GO) Term Enrichment analysis, chord diagrams, and multiple comparisons were generated using iPathwayGuide™ (AdvaitaBio Corporation, https://ipathwayguide.advaitabio.com/)^39,40^. p-values were adjusted for multiple comparisons controlling false discovery rate (FDR) using the Benjamini-Hochberg (BH) procedure^41^. Significant GO terms in the stimulated vs unstimulated homeRNA group (Fig 2B) were selected at an FDR<0.05. To perform multiple comparisons and determine overlapping GO biological processes between all three experimental stimulated vs unstimulated conditions seen in Figure 3B and 3F, the high specificity pruning method was used to assess the significance of GO terms. KEGG pathways analysis was also performed using iPathwayGuide™ (AdvaitaBio Corporation, https://ipathwayguide.advaitabio.com/)^39,40^. Significant KEGG pathways between the stimulated vs unstimulated conditions across all three groups seen in Figure 3C and 3G were selected using an FDR<0.05.

**Figure 1.**
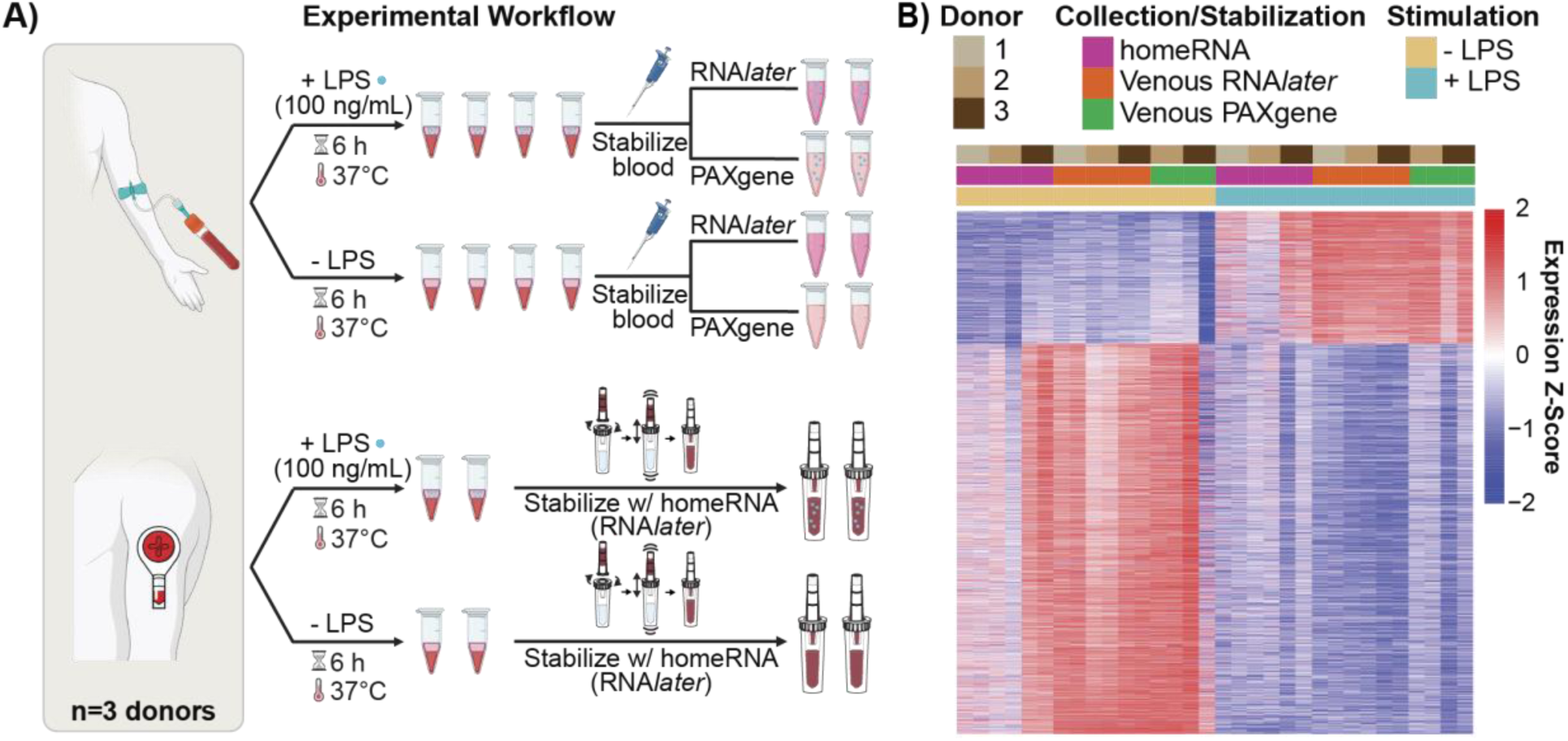
Comparison of response to lipopolysaccharide (LPS) in venous blood and homeRNA-stabilized blood. A) Workflow of stimulating blood with LPS in venous blood and Tasso-collected blood. Each donor (n=3 donors) had blood drawn from a phlebotomy draw and from the upper arm using the Tasso-SST device on the same day. The blood samples were then aliquoted and incubated in media supplemented with 100 ng/mL LPS or in control media (without LPS). After incubation, blood samples collected from the Tasso device were then stabilized with homeRNA, and blood samples collected from the phlebotomy draw were stabilized with either RNA*later* or PAXgene. Created using Biorender.com. B) Unsupervised heatmap of differentially expressed genes (30% change in expression, FDR ≤ 0.05) between LPS-stimulated and unstimulated blood samples across different collection and stabilization methods after removing the batch effect from different donors. Each column represents an individual sample (each donor had two replicates), and each row represents a gene (7777 genes in total). The homeRNA and Venous RNA*later* samples had three donors (n=6 samples) and Venous PAXgene had two donors (n=4 samples). Colors represent standardized expression values (z-scores) for each gene across all samples, with red indicating higher expression and blue indicating lower expression relative to each gene’s mean across all experimental conditions.

**Figure 2.**
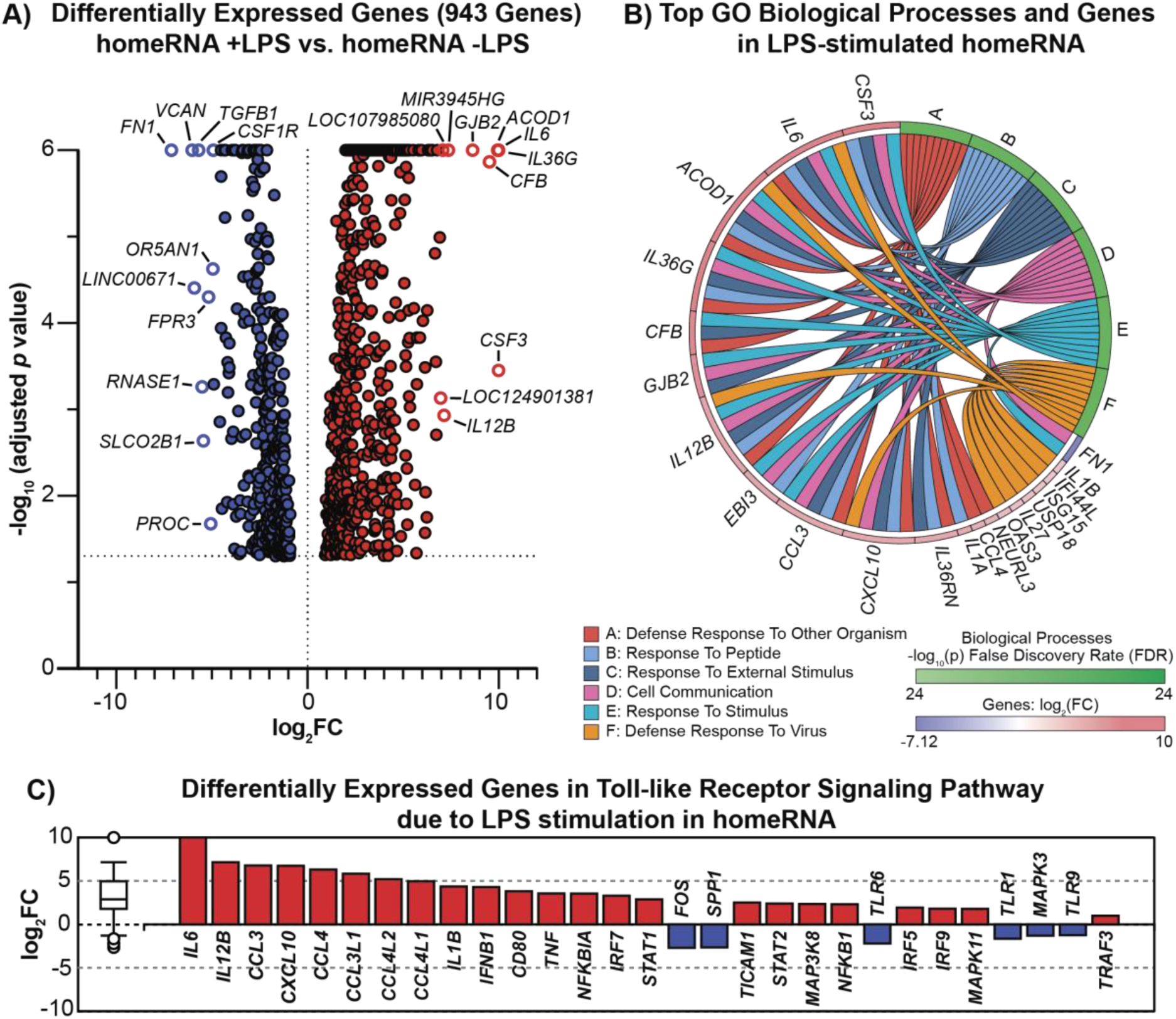
homeRNA captures LPS-induced immune response. A) Volcano plot of differentially expressed genes (DEGs) between LPS-stimulated and unstimulated homeRNA-stabilized samples. The measured 943 DEGs (30% change in expression, FDR ≤ 0.05) are represented by the negative log_10_ of the adjusted p-value on the y-axis and log_2_ fold change on the x-axis. Red points represent up-regulated genes and blue points represent down-regulated genes with LPS stimulation. The top 10 highly expressed up-regulated and down-regulated genes are labeled. B) Gene ontology (GO) enrichment chord diagram visualizing the top 10 DEGs present in each of the top six biological processes. Genes are positioned around the circle and colored by their log_2_ fold change (red for up-regulated, blue for down-regulated). Biological processes are labeled A-F around the outer ring. Connecting ribbons link genes to their associated biological processes, with ribbon colors corresponding to a biological process. C) All up- and down-regulated DEGs (n=29 DEGs) in the toll-like receptor signaling pathway (KEGG: 04060) ranked based on their absolute value of log_2_ fold change. Up-regulated genes are shown in red; down-regulated genes are shown in blue. The box and whisker plot on the left summarizes the distribution of all DEGs in this pathway with outliers represented by circles. p-values were adjusted for multiple comparisons by controlling for the false discovery rate (FDR) using the Benjamini-Hochberg (BH) procedure^41^.

**Figure 3.**
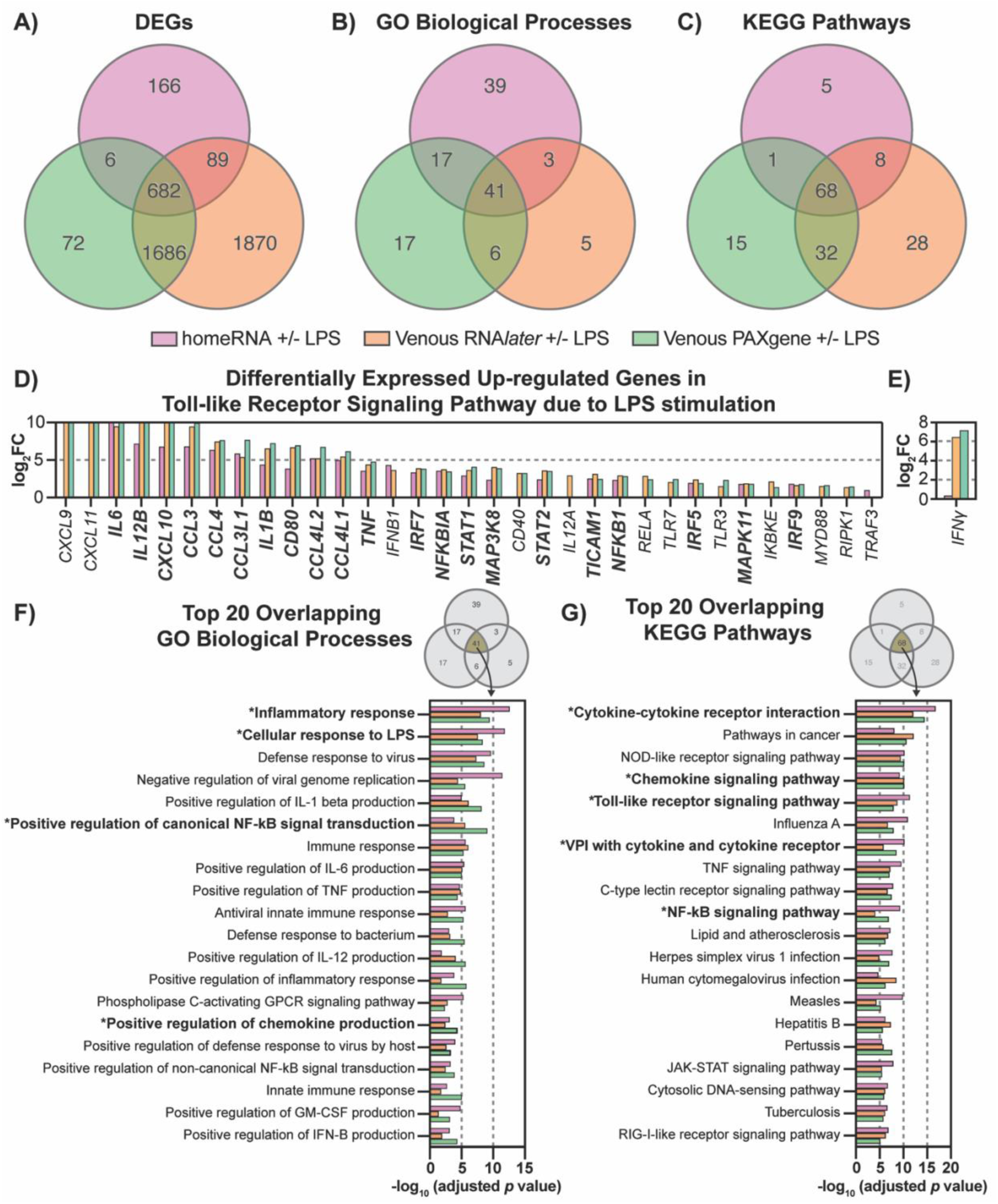
homeRNA-stabilized samples capture a similar LPS-induced inflammatory response to that of venous blood stabilized with RNA*later* or PAXgene. Overlapping A) differentially expressed genes (DEGs) between the LPS stimulated and unstimulated conditions of the homeRNA-stabilized, RNA*later*-stabilized venous blood, and PAXgene-stabilized venous blood samples. B) Enriched gene ontology (GO)-derived biological processes and C) KEGG pathways from the DEGs. D) All up-regulated DEGs (n=33 DEGs) in the toll-like receptor signaling pathway (KEGG: 04060) from each LPS-stimulated and unstimulated comparison. DEGs are ranked based on their average log_2_ fold change across all three groups. E) LPS-stimulated response of *IFNγ* in each comparison. F) Top 20 overlapping GO-derived biological processes pathways between all three LPS-stimulated collection/stabilization groups. Biological processes are ranked based on their average adjusted p-value across all three groups. G) Top 20 overlapping KEGG pathways between all three LPS-stimulated collection/stabilization groups. Pathways are ranked based on their average adjusted p-value across all three groups. VPI stands for viral protein interaction. p-values were adjusted for multiple comparisons by controlling for the false discovery rate (FDR) using the Benjamini-Hochberg (BH) procedure^41^.

## RESULTS & DISCUSSION

### Comparison of LPS-induced gene expression across homeRNA and venous blood collection methods

In this study, we sought to assess the ability of the homeRNA kit to capture a biological response through bulk RNA-sequencing methods. To accomplish this, we induced a known biological response via bacterial lipopolysaccharide (LPS) stimulation, a well established immune *ex vivo* whole blood stimulation model that activates a targeted inflammatory response^31–33^. To further establish homeRNA as a remote sampling tool that can be used to probe transcriptomic immune responses, we also compared the LPS-stimulated response of homeRNA-stabilized samples to that of RNA*later*-stabilized or PAXgene-stabilized venous blood. These stabilizers were chosen as RNA*later* is the stabilizer used in homeRNA, allowing for direct comparison of homeRNA-collected and stabilized samples with phlebotomy drawn and stabilized blood. PAXgene was chosen as it is a commonly used stabilization solution for clinic-based phlebotomy draws since it comes already packaged in vacutainer tubes. To properly compare these different collection and stabilization methods with a LPS-stimulated response, we designed an experiment where venous blood (via phlebotomy draw) and capillary blood (using the lancet-based Tasso-SST included in the homeRNA kit that draws blood from the upper arm) were collected from a participant during the same visit. All collected blood samples (Tasso-collected blood and phlebotomy-drawn blood) were either stimulated with 100 ng/mL LPS or left unstimulated prior to stabilization (Figure 1A). Venous blood samples were stabilized with either RNA*later* or PAXgene (Figure 1A). Tasso-collected blood samples were stabilized by attaching the Tasso tube to a custom engineered stabilizer tube containing RNA*later* and shaking per the instructions of the homeRNA kit (Figure 1A)^6,27–30^.

First, to illustrate the overall gene expression response of the three different collection and stabilization methods, we generated a heatmap of differentially expressed genes associated with response to LPS (Figure 1B). The heatmap reveals that all three collection/stabilization methods captured the expected changes in gene expression following LPS stimulation, with clear differential up-regulation and down-regulation of genes in response to LPS. However, some donor-to-donor variability is evident, particularly within donor 3 in the homeRNA and Venous PAXgene samples (Figure 1B). This variability likely reflects a combination of biological differences in individual immune responses to LPS stimulation, technical variations inherent to each collection and stabilization protocol, and in the case of homeRNA, potential differences between capillary and venous blood composition. Despite this donor variability, a differential LPS-induced response is still observed across the different collection and stabilization methods.

### homeRNA captured LPS-induced biological response

We next interrogated the ability of homeRNA to capture an induced biological response via LPS stimulation. We identified 943 differentially expressed genes (DEGs) with LPS stimulation in the homeRNA-stabilized samples; the complete list of DEGs and significantly expressed KEGG pathways between the LPS-stimulated and unstimulated homeRNA-stabilized samples are shown in Supplementary Table S3. Figure 2A summarizes the identified DEGs, with the top 10 most significantly up- and down-regulated genes in response to LPS stimulation in Tasso-collected and homeRNA-stabilized blood samples labeled. A number of these genes are associated with inflammatory responses and have been shown previously to be expressed in LPS-stimulated blood. These genes include: *IL6* and *IL12B*, both of which are cytokines activated downstream of LPS-induced toll-like receptor 4 (TLR4) pathway^42,43^; *IL36G* and *CSF3*, cytokines present in regulating innate inflammatory responses mediated by IL-1 and IL-6, respectively^44–47^; and *CFB* and *ACOD1*, which are regulators in LPS-activated immune response pathways^48,49^.

Further, gene ontology enrichment analysis was performed to identify the top six biological processes and the top 10 overrepresented genes within each process (Figure 2B). These top gene ontology (GO)-derived biological processes are related to defense responses to pathogens, peptides, and other stimuli. Overrepresentation of these biological processes is expected as LPS is a molecule present on the surface of gram-negative bacteria and thus stimulates a host-like response. Notably, cytokines (*IL6*, *IL36G*, *IL12B*), complement components (*CFB*), and chemokines (*CCL3*, *CXCL10*) were strongly linked to immune defense pathways, suggesting an overall innate immune activation due to LPS.

Lastly, we further interrogated the alterations in gene expression related to the toll-like receptor signaling (Figure 2C). Toll-like receptor signaling is the primary pathway activated in an LPS response; LPS binds TLR4 in immune cells, transducing signals across the plasma membrane^50^. TLR4 activation then initiates key downstream inflammatory pathways including mitogen-activated protein kinase (MAPK) signalling and nuclear factor kappa (NF-kB) signalling, resulting in an inflammatory response^51–53^. When we probed the toll-like receptor signaling pathway, we observed an upregulation in cytokines (*IL6*, *IL12B*, *IL1B*, and *TNF*), chemokines (*CXCL10*, *CCL3*, *CCL3L1*, *CCL4*, *CCL4L1*, *CCL4L2*), and interferon related genes (*IFNB1*, *IRF7*) that is consistent in LPS-induced immune response in the literature (Figure 2C)^32,42,54–56^. Similarly, we observed similar LPS-induction of genes associated with the NF-kB and MAPK signaling pathways, such as *IL1A*, *IL1B*, and *ICAM1*, demonstrating the full inflammatory response captured in LPS-induced, homeRNA-stabilized samples (Supplementary Figure S2).

### homeRNA-stabilized samples capture a similar LPS-induced inflammatory response to that of venous blood stabilized with RNAlater or PAXgene

Having demonstrated that homeRNA can effectively capture biological responses via LPS stimulation, we next sought to directly compare the induced immune response in homeRNA-stabilized capillary blood with that of traditional phlebotomy-drawn venous blood. The primary aim of this comparative analysis was to characterize differences in gene expression profiles between these two collection methods to inform the feasibility of using homeRNA in future transcriptomic studies. This comparison is particularly relevant for determining whether homeRNA could enable the transition of clinic-based immune response studies to remote sampling applications. By comparing the similarities and differences between homeRNA (capillary) and venous blood transcriptomic responses to LPS stimulation, we aim to provide critical guidance for researchers considering the implementation of homeRNA-based remote blood collection in studies investigating immune activation and inflammatory pathways.

Towards this aim, we first identified DEGs in LPS-stimulated and unstimulated blood samples within our three collection/stabilization methods: Tasso-collected blood stabilized with RNA*later* (i.e., homeRNA-stabilized samples), venous blood stabilized with RNA*later*, and venous blood stabilized with PAXgene. The complete list of DEGs and significantly expressed KEGG pathways for the LPS-stimulated and unstimulated comparisons of the homeRNA-stabilized, Venous RNA*later*, and Venous PAXgene groups are shown in Supplementary Tables S3, S4, and S5, respectively. We found 682 DEGs to be common between all three groups (Figure 3A), suggesting some alignment between the LPS-induced response from each collection and stabilization method. In addition, each method captured unique DEGs, suggesting that some components of the response may be method-specific (i.e., collection method or stabilization method), potentially reflecting differences in sample type (i.e., capillary vs. venous). GO-derived biological processes and KEGG pathway analysis on the DEGs of each group also found 41 overlapping biological processes (Figure 3B) and 68 overlapping pathways (Figure 3C); the complete list of overlapping biological processes and pathways between the LPS-stimulated and unstimulated conditions of the three groups are shown in Supplementary Tables S6 and S7, respectively.

To examine these differences in more detail, we focused on the toll-like receptor signaling pathway, a key mediator of LPS-induced immune responses. We identified 33 up-regulated DEGs in the toll-like receptor signaling pathway induced by LPS stimulation across the three collection/stabilization groups (Figure 3D). The majority of these up-regulated DEGs were consistently present in all three groups, including key LPS-induced cytokines (*IL6*, *IL12B*, *IL1B*, *TNF*) and chemokines (*CXCL10*, *CCL3*, *CCL4*) as previously described. However, we observed a notable difference in the expression of the two chemokines *CXCL9* and *CXCL11*, which showed significant up-regulation in both venous blood groups (RNA*later* and PAXgene) following LPS stimulation, but were absent from the stimulated homeRNA blood samples (Figure 3D). This difference appears to be linked to interferon-gamma (IFN-γ) signaling, as CXCL9 and CXCL11 are specifically IFN-γ-inducible chemokines while CXCL10, which is present in all three groups, can be induced by LPS or TNF-ɑ as well as IFN-γ^57–59^. Consistent with this IFN-γ signaling, we observed significant up-regulation of *IFNγ* in both LPS-stimulated venous groups while no significant change was observed in stimulated homeRNA samples (Figure 3E). The absence of *IFNγ* induction in homeRNA samples may explain why *CXCL9* and *CXCL11* were not differentially expressed in these samples. This is further supported by previous studies demonstrating that LPS and IFN-γ act synergistically to induce CXCL9 and CXCL11 expression in various cell types^60–63^, providing a potential mechanistic basis for the robust up-regulation of these chemokines observed specifically in the venous blood samples where both LPS stimulation and *IFNγ* induction occurred. It should be noted that other studies found that LPS stimulation did not induce IFN-γ protein in either capillary blood collected via a microneedle in the upper arm or venous blood^64^, but yet other studies have found that LPS can induce IFN-γ in natural killer cells and monocytes^65,66^. Taken together, there is variation observed in the literature in LPS-induced IFN-γ levels across blood types; the observed differences in transcriptional *IFNγ* induction between capillary and venous blood in our study could be a result of physiological differences between capillary and venous blood. Given our limited sample size, future studies with larger cohorts would be needed to provide greater statistical power and more definitive insights into IFN-γ-mediated differences between these sampling methods.

Despite differences in *IFNγ* induction and baseline inflammatory signature between the homeRNA and RNA*later*-stabilized venous samples, we were still able to capture a similar inflammatory response to LPS stimulation across all blood collection and stabilization conditions. These similarities in LPS response are further seen in the top 20 overlapping overrepresented biological processes (Figure 3F) and pathways (Figure 3G) across all three groups. Here, as expected, we observe significant enrichment of biological processes and pathways related to inflammation and host immune response across all groups due to LPS stimulation. Overall, the primary LPS-driven immune response signature remains well-aligned across the collection and stabilization methods, suggesting that homeRNA can capture a similar inflammatory response to that of venous blood sampling.

We also compared the baseline gene expression profiles between unstimulated homeRNA-stabilized blood (capillary) and RNA*later*-stabilized venous blood (Supplementary Figures S3, S4, and S5 and associated discussion). We identified a baseline inflammatory signature present in capillary samples and absent in venous samples that likely reflects methodological differences in blood collection completed in this study; specifically, the Tasso-SST blood collection tubes used for the homeRNA samples contain no anticoagulant, whereas the venous blood samples were collected in EDTA-coated vacutainers. The lack of anticoagulant in the Tasso-collected samples may have promoted clotting activation, contributing to the observed inflammatory signature. Additionally, in this study the collected blood samples were incubated for six hours prior to stabilization so that they could serve as controls for LPS-stimulated samples, which were also incubated for six hours. However, in typical homeRNA-based studies, collected capillary blood samples would be immediately stabilized by the participant after collection. Therefore, the increased gene expression response observed in some inflammatory genes between homeRNA and RNA*later*-stabilized venous samples in this study does not necessarily translate to typical conditions used in homeRNA studies. Despite some of the inflammatory genes being up-regulated in homeRNA-stabilized, unstimulated samples, when we probed the bulk gene expression profile (19,751 genes total), genome-wide analysis across all donors revealed strong concordance between homeRNA and venous RNA*later* samples, with a group-level correlation of r=0.948 (Pearson correlation r=0.948) (Supplementary Figure S3). This correlation is consistent with other comparisons on the gene and protein expression between capillary blood and venous blood observed in other studies^68–72^, supporting the potential of homeRNA to be used in remote transcriptomic studies in place of clinic-based phlebotomy draws.

To further assess the similarity of transcriptomic profiles between collection and stabilization methods at the individual sample level, we performed pairwise correlation analyses of gene expression across all samples for each donor (Figure 4 for Donor 1; Supplementary Figures S6 and S7 for Donors 2 and 3). These donor-specific analyses revealed high technical reproducibility within each collection and stabilization method, with most correlations between technical replicates exceeding r = 0.92 across all conditions and donors (range: r = 0.928 to 0.957 for homeRNA; r = 0.951 to 0.956 for venous RNA*later*; r = 0.883 to 0.951 for venous PAXgene). Between-method comparisons at the sample level also demonstrated strong correlations under both unstimulated and LPS-stimulated conditions across all three donors. For unstimulated samples, correlations between homeRNA and venous RNA*later* ranged from r = 0.886 to 0.922, while correlations between homeRNA and venous PAXgene ranged from r = 0.829 to 0.903 (Donors 2 and 3 only; Supplementary Figure S6-7). For LPS-stimulated samples, correlations between homeRNA and venous RNAlater ranged from r = 0.861 to 0.902, while correlations between homeRNA and venous PAXgene ranged from r = 0.852 to 0.890. Notably, the two venous collection methods (RNA*later* and PAXgene stabilization) also showed high correlation with each other at the sample level (range: r = 0.857 to 0.930 for the LPS-stimulated and unstimulated conditions; Supplementary Figures S6-7), indicating that differences in stabilization reagent have minimal effects on overall transcriptomic profiles. These consistently high correlations across donors, conditions, and methods, both at the group level (r = 0.948 for homeRNA vs. venous RNA*later* and venous RNA*later* vs. venous PAXgene; Supplementary Figures S3-4) and individual sample level (full range r = 0.856-0.930 for all between-method comparisons; Figure 4, Supplementary Figures S6-7), demonstrate that homeRNA-stabilized capillary blood produces LPS-induced transcriptomic profiles that closely mirror those obtained from traditional venous blood collection. Overall, these results support the use of homeRNA as a viable alternative for genome-wide transcriptomic studies requiring remote blood collection.

**Figure 4.**
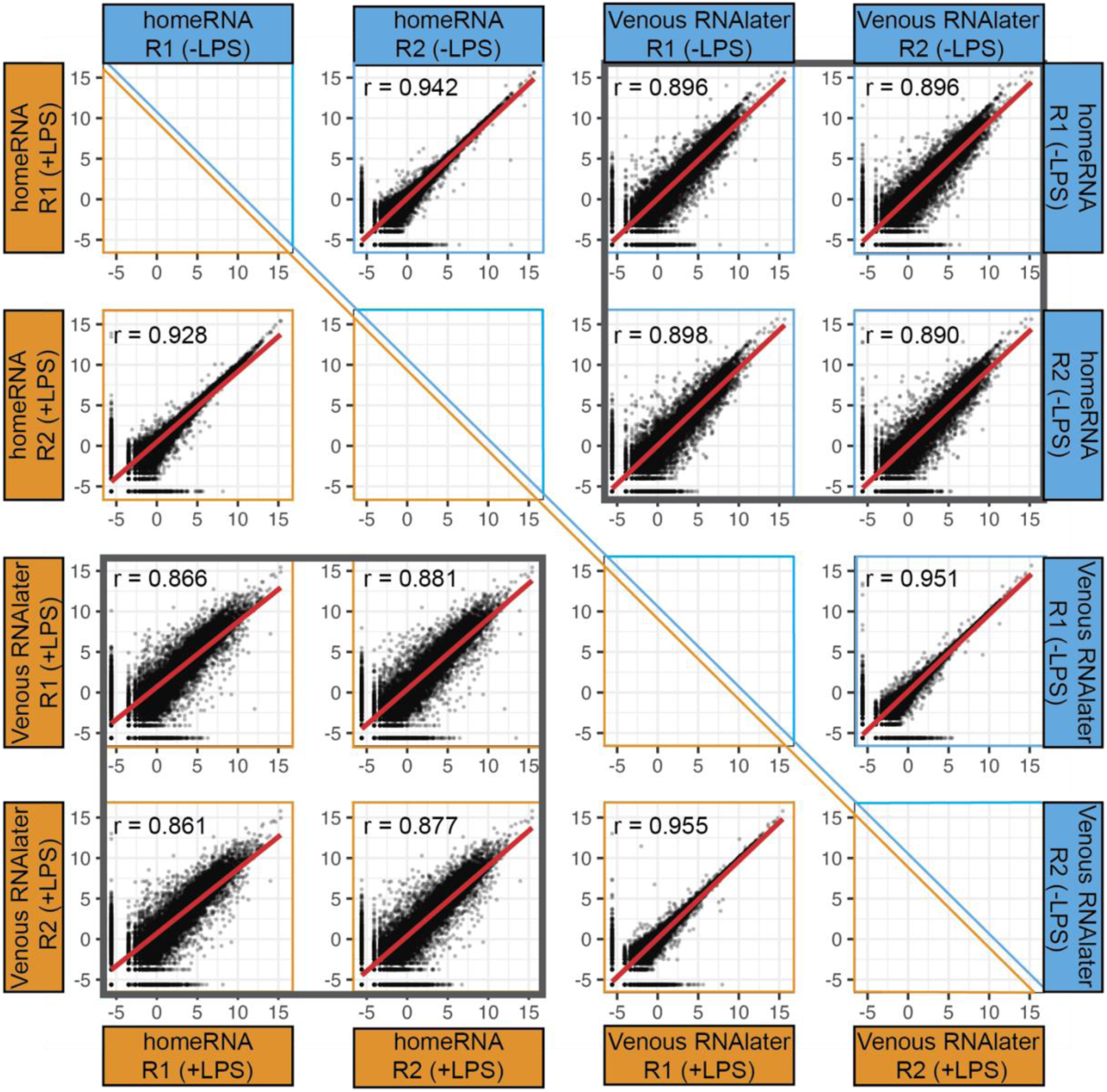
LPS-stimulated (orange) and unstimulated (blue) gene expression profiles are similar between homeRNA-stabilized capillary blood and RNA*later*-stabilized venous blood samples in Donor 1. Scatter plots compare log2-transformed normalized gene expression values (logCPM) between replicates of homeRNA capillary blood (Tasso-collected and RNA*later*-stabilized) samples and RNA*later*-stabilized venous blood samples. Unstimulated samples (-LPS) are plotted in the upper right triangle (blue panels) and LPS-stimulated samples (+LPS) are plotted in the lower left triangle (orange panels). Each sample was collected and processed in technical replicates (R1 and R2). Within-method comparisons show high technical reproducibility between replicates for both homeRNA (r = 0.942 for -LPS; r = 0.928 for +LPS) and Venous RNA*later* (r = 0.951 for -LPS; r = 0.955 for +LPS). Between-method comparisons (boxed in gray) show strong correlation between homeRNA and venous RNA*later* samples under both unstimulated (r = 0.896 to 0.898) and LPS-stimulated (r = 0.861 to 0.881) conditions, demonstrating that homeRNA-stabilized capillary blood produces transcriptomic profiles highly similar to traditional venous blood collection. Red lines indicate the line of best fit. Pearson correlation coefficients (r) are displayed in each panel.

### Limitations

Several limitations of this study should be noted. First, our sample size (n=4-6 samples from 2-3 unique donors for each group) limits our statistical power to detect subtle differences and increases the potential impact of donor variability. The differences we observed in LPS-induced *IFNγ* expression, for example, could reflect true biological or technical differences, or could be influenced by donor-to-donor variability amplified by small sample size. The literature on LPS-induced IFN-γ production is inconsistent; some studies report no protein IFN-γ induction in either capillary or venous blood^65^, while others demonstrate IFN-γ production in specific immune cell subsets^66,67^. This differential response could reflect biological differences in immune cell composition in capillary vs. venous blood, technical differences related to anticoagulant presence affecting cellular responses (the Tasso-SST tube lacks an anticoagulant while venous blood is collected in EDTA-coated vacutainers), or donor-to-donor variability. Distinguishing between these possibilities would require controlled experiments comparing capillary blood collected with and without anticoagulants, as well as larger sample sizes to account for donor variability.

Second, our experimental design incorporated a 6-hour incubation period prior to stabilization for both stimulated and unstimulated samples, which differs from standard homeRNA protocols that call for immediate stabilization. This extended incubation period of blood collected in the Tasso-SST tube (which contains no anticoagulant) likely amplified the baseline inflammatory signature in homeRNA samples compared to the venous blood samples that were collected in EDTA-coated tubes and incubated. Specifically, the absence of anticoagulant in Tasso-SST tubes would be expected to initiate clotting cascades and activate inflammatory pathways, contributing to the baseline inflammatory signature observed in homeRNA samples. Therefore, the baseline gene expression profile of unstimulated homeRNA samples found in this study may not reflect the baseline gene expression profiles that would be observed with immediate stabilization typical in remote use of homeRNA.

Third, our study design confounds multiple variables, mainly blood source (capillary vs. venous), collection method (Tasso device vs. phlebotomy draw), stabilization efficiency (shaking in custom tube vs. pipetting/inversion in standard tubes), and anticoagulant presence, making it difficult to definitively attribute observed differences to any single factor. Future studies with larger sample sizes, systematic variation of individual parameters, and immediate stabilization protocols would help address these limitations and provide more definitive guidance on the sources of variation between collection methods.

## CONCLUSION

In this study, we demonstrate the feasibility of homeRNA for transcriptomic studies by comparing the gene expression profile and LPS-induced biological response between homeRNA sampling and traditional venous sampling. We found that homeRNA successfully captured a LPS-induced inflammatory response that was comparable to that of venous blood samples stabilized with either RNA*later* or PAXgene (682 shared DEGs, 41 shared biological processes, 68 shared KEGG pathways, and consistent up-regulation of key inflammatory mediators including *IL6*, *IL1B*, *TNF*, and multiple chemokines). Further, we observed strong correlation in baseline gene expression profiles between the two collection methods (capillary blood from the upper arm and venous blood) and the two stabilization methods in venous blood (RNA*later* and PAXgene). This work establishes the compatibility of homeRNA with total RNA-sequencing, demonstrating its potential as a useful tool for monitoring of whole-transcriptome immune response via remote sampling.

Overall, this study aims to provide guidance for other researchers seeking to combine remote blood collection studies with downstream RNA-sequencing analysis. Future work with larger sample sizes will be required to further validate the biological reproducibility of homeRNA and to establish which genes and pathways can most accurately be quantified using homeRNA. To date, we have already demonstrated the use of homeRNA to capture dynamic immune response signatures during SARS-CoV-2 infection, including both the presymptomatic and acute phases, and immune response to wildfire smoke exposure via a targeted gene panel method^6,30,31^. This current work adds to the body of literature that characterizes the biological and analytical factors that contribute to transcriptomic differences between capillary and venous blood sampling that is crucial to properly interpret results in remote, capillary-blood based studies compared to clinic-based venipuncture studies.

## Supporting information

Aligned Gene Counts for All Samples

## ACKNOWLEDGEMENTS

This publication was supported by Schmidt Sciences, LLC, the David and Lucile Packard Foundation (for device manufacturing and human subject components of the research), and the National Institutes of Health (NIH) through the National Institute of General Medical Sciences award number R35GM128648, the National Center for Advancing Translational Sciences award number 5TL1TR002318-08 (for support of LGB), and the University of Washington EDGE Center of the National Institute of Health award number P30ES007033 (for support of JMW and TKB and for pathway analysis software). This publication was made possible through funding from the Fulbright Program (for support of LAM). The REDCap used for human subjects consent and enrollment is supported by the Institute of Translational Health Sciences, which is funded by the National Center for Advancing Translational Sciences of the National Institutes of Health under award number UL1TR002319. The content is solely the responsibility of the authors and does not necessarily represent the official views of the National Institutes of Health or other funding bodies. We would like to thank Sean Bennett for helpful discussions on RNA-sequencing and analysis. We would also like to thank the study participants.

