## Supplementary Information for "From Home to Transcriptome: Comparing the transcriptomic profile of induced immune response via lipopolysaccharide stimulation in homeRNA and venous blood"

**Table of Contents:**

| **Supplemental Tables** | S2-3 |
| --- | --- |
| **Supplemental Table Descriptions** | S4 |
| **Supplemental Figures** | S5-6, S8-12 |
| **Supplemental Discussion of Figures S2-4** | S7 |
| **References** | S13 |

**TABLES**

**Table S1**: Individual Sample Read Count

| **Sample** | **Donor** | **Replicate** | **Was the sample stimulated with LPS?** | **Total reads (millions)** | **Aligned reads (millions)** | **Alignment Percentage (%)** |
| --- | --- | --- | --- | --- | --- | --- |
| homeRNA (Tasso RNA*later*) | 1 | 1 | yes | 51.06 | 19.00 | 37.2 |
| homeRNA (Tasso RNA*later*) | 1 | 2 | yes | 76.62 | 33.74 | 44.0 |
| homeRNA (Tasso RNA*later*) | 2 | 1 | yes | 54.44 | 23.86 | 43.8 |
| homeRNA (Tasso RNA*later*) | 2 | 2 | yes | 54.46 | 23.21 | 42.6 |
| homeRNA (Tasso RNA*later*) | 3 | 1 | yes | 60.98 | 26.29 | 43.1 |
| homeRNA (Tasso RNA*later*) | 3 | 2 | yes | 63.24 | 25.53 | 40.4 |
| homeRNA (Tasso RNA*later*) | 1 | 1 | no | 62.47 | 28.99 | 46.4 |
| homeRNA (Tasso RNA*later*) | 1 | 2 | no | 67.79 | 30.74 | 45.4 |
| homeRNA (Tasso RNA*later*) | 2 | 1 | no | 68.58 | 27.37 | 39.9 |
| homeRNA (Tasso RNA*later*) | 2 | 2 | no | 57.37 | 24.48 | 42.7 |
| homeRNA (Tasso RNA*later*) | 3 | 1 | no | 69.38 | 28.58 | 41.2 |
| homeRNA (Tasso RNA*later*) | 3 | 2 | no | 98.27 | 38.16 | 38.8 |
| Venous PAXgene | 2 | 1 | yes | 53.41 | 17.10 | 32.0 |
| Venous PAXgene | 2 | 2 | yes | 54.29 | 17.33 | 31.9 |
| Venous PAXgene | 3 | 1 | yes | 50.97 | 19.19 | 37.7 |
| Venous PAXgene | 3 | 2 | yes | 54.13 | 18.87 | 34.9 |
| Venous PAXgene | 2 | 1 | no | 54.01 | 16.00 | 29.6 |
| Venous PAXgene | 2 | 2 | no | 56.73 | 16.47 | 29.0 |
| Venous PAXgene | 3 | 1 | no | 53.88 | 16.22 | 30.1 |
| Venous PAXgene | 3 | 2 | no | 55.29 | 20.42 | 36.9 |
| Venous RNA*later* | 1 | 1 | yes | 72.95 | 33.34 | 45.7 |
| Venous RNA*later* | 1 | 2 | yes | 56.39 | 24.27 | 43.0 |
| Venous RNA*later* | 2 | 1 | yes | 73.93 | 30.62 | 41.4 |
| Venous RNA*later* | 2 | 2 | yes | 81.14 | 33.96 | 41.9 |
| Venous RNA*later* | 3 | 1 | yes | 50.84 | 21.65 | 42.6 |
| Venous RNA*later* | 3 | 2 | yes | 51.07 | 21.10 | 41.3 |
| Venous RNA*later* | 1 | 1 | no | 50.78 | 21.69 | 42.7 |
| Venous RNA*later* | 1 | 2 | no | 51.12 | 21.46 | 42.0 |
| Venous RNA*later* | 2 | 1 | no | 67.19 | 26.42 | 39.3 |
| Venous RNA*later* | 2 | 2 | no | 52.64 | 22.18 | 42.1 |
| Venous RNA*later* | 3 | 1 | no | 56.99 | 24.42 | 42.9 |
| Venous RNA*later* | 3 | 2 | no | 54.77 | 22.89 | 41.8 |

**Table S2**: Summary Read Count of Blood Collection/Stabilization Groups

| **Collection/Stabilization Method** | **# of Samples** | **Total reads (millions)** | **Aligned reads (millions)** | **Alignment Percentage (%)** |
| --- | --- | --- | --- | --- |
| homeRNA (Tasso RNA*later*) | 12 | 65.4 ± 12.7 | 27.5 ± 5.1 | 42.1 ± 2.7 |
| Venous RNA*later* | 12 | 60.0 ± 10.8 | 25.3 ± 4.7 | 42.2 ± 1.5 |
| Venous PAXgene | 8 | 54.1 ± 1.6 | 17.7 ± 1.6 | 32.8 ± 3.3 |

**Table Descriptions (separate table files not included in SI):**

**Supplementary Table 3 (xslx):** LPS-stimulated transcriptomic response compared to unstimulated response in homeRNA-collected and stabilized samples. This table lists all up-regulated and down-regulated differentially expressed genes (DEGs) (|log2FC| ≥ 1.3, adjusted p-value ≤ 0.05), significantly enriched GO-derived biological processes (FDR < 0.05), and significant KEGG pathways (FDR < 0.05) present in the comparison of LPS-stimulated homeRNA samples to unstimulated homeRNA samples.

**Supplementary Table 4 (xslx):** LPS-stimulated transcriptomic response compared to unstimulated response in RNAlater-stabilized venous blood samples. This table lists all up-regulated and down-regulated differentially expressed genes (DEGs) (|log2FC| ≥ 1.3, adjusted p-value ≤ 0.05), significantly enriched GO-derived biological processes (FDR < 0.05), and significant KEGG pathways (FDR < 0.05) present in the comparison of LPS-stimulated RNAlater-stabilized venous blood samples to unstimulated RNAlater-stabilized venous blood samples.

**Supplementary Table 5 (xslx):** LPS-stimulated transcriptomic response compared to unstimulated response in PAXgene-stabilized venous blood samples. This table lists all up-regulated and down-regulated differentially expressed genes (DEGs) (|log2FC| ≥ 1.3, adjusted p-value ≤ 0.05), significantly enriched GO-derived biological processes (FDR < 0.05), and significant KEGG pathways (FDR < 0.05) present in the comparison of LPS-stimulated PAXgene-stabilized venous blood samples to unstimulated PAXgene-stabilized venous blood samples.

**Supplementary Table 6 (xslx):** Complete list of enriched GO-derived biological processes with high-specificity pruning p-value correction (p-elim < 0.05) between all three LPS-stimulated collection/stabilization groups. This table lists the intersecting and unique biological processes present in the LPS-stimulated and unstimulated comparisons of the homeRNA-stabilized capillary, RNAlater-stabilized venous, and PAXgene-stabilized venous samples.

**Supplementary Table 7 (xslx):** Complete list of significant KEGG pathways (FDR < 0.05) between all three LPS-stimulated collection/stabilization groups. This table lists the intersecting and unique pathways present in the LPS-stimulated and unstimulated comparisons of the homeRNA-stabilized capillary, RNAlater-stabilized venous, and PAXgene-stabilized venous samples.

**FIGURES**

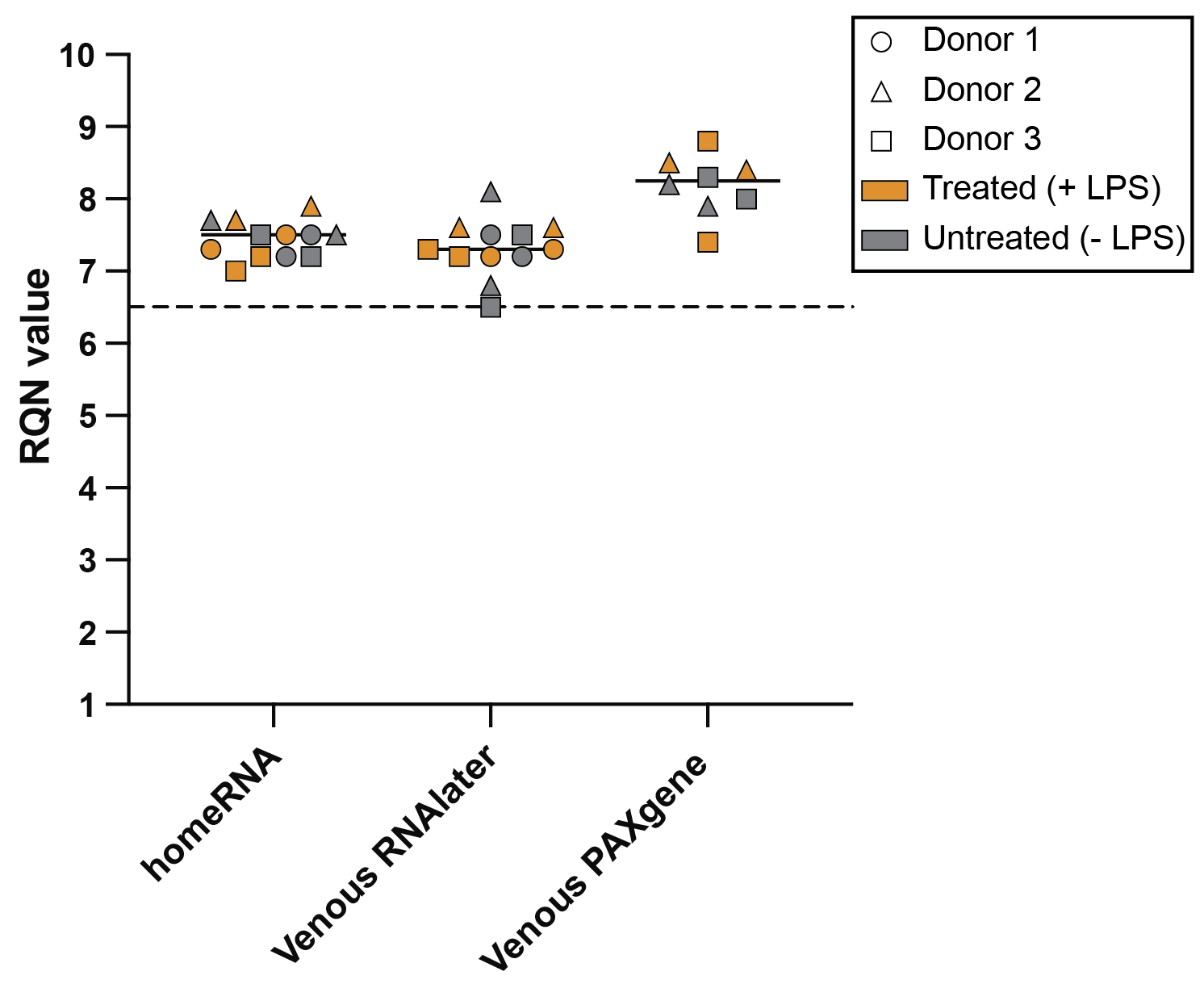

**Figure S1**. RNA Quality Number (RQN) distributions of isolated RNA from all blood collection and stabilization methods prior to library preparation and total RNA-sequencing. Each point represents an individual isolated RNA sample. Circles, triangles, and squares represent RNA samples from Donor 1, Donor 2, and Donor 3, respectively. Samples that were treated with 100 ng/mL lipopolysaccharide (LPS) are colored in orange and samples that were untreated are colored in gray. RQN values range from 1 to 10, with 1 representing completely degraded RNA and 10 representing completely intact RNA. All samples from this study had RQN values ≥ 6.5, with homeRNA-collected and -stabilized capillary blood, RNA*later*-stabilized venous blood, and PAXgene-stabilized venous blood samples having an average RQN of 7.4, 7.3, and 8.2, respectively.

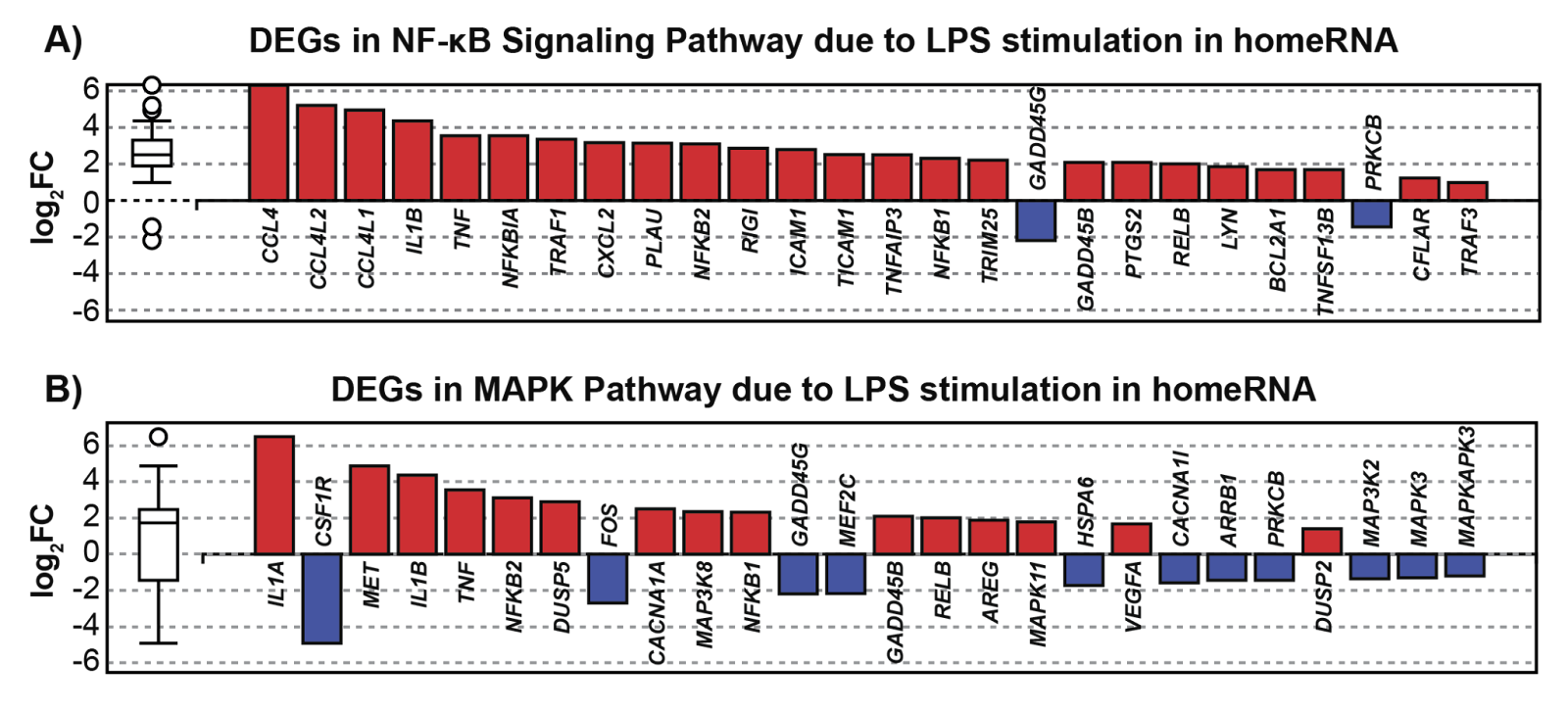
 **Figure S2**. Differentially expressed genes (DEGs) in downstream MAPK and NF-kB pathways initiated by LPS stimulation in homeRNA-collected and stabilized samples. A) All up- and down-regulated DEGs (n=26 DEGs) in the NF-kB signaling pathway (KEGG: 04064) ranked based on their absolute value of log2 fold change. B) All up- and down-regulated DEGs (n=26 DEGs) in the mitogen-activated protein kinase (MAPK) pathway (KEGG: 04010) ranked based on their absolute value of log2 fold change. Up-regulated genes are shown in red; down-regulated genes are shown in blue. The box and whisker plot on the left summarizes the distribution of all DEGs in this pathway with outliers represented by circles.

***Supplementary discussion of Figures S3, S4, and S5***

In addition to LPS stimulation, we compared the baseline gene expression profiles between unstimulated homeRNA-stabilized blood (capillary) and unstimulated venous blood stabilized with RNA*later* (the same stabilization reagent used in homeRNA). We identified 1489 DEGs between unstimulated capillary and venous blood samples (Figure S3A), with many of the up-regulated genes in the capillary samples being cytokines and chemokines associated with inflammatory responses (e.g., *CCL2*, *CXCL2*, *CXCL3*, and *IL1A*). This baseline inflammatory signature in capillary samples likely reflects methodological differences in blood collection. Specifically, the Tasso-SST blood collection tubes used for the homeRNA samples contain no anticoagulant, whereas the venous blood samples were collected in EDTA-coated vacutainers. Consequently, we observed visible coagulation in the Tasso-collected capillary samples immediately following blood collection but not in the venous samples. This coagulation process may have triggered a baseline inflammatory response in the capillary samples, accounting for the elevated expression of inflammatory mediators even without LPS stimulation.

Importantly, in this study the collected blood samples were incubated for six hours prior to stabilization so that they could serve as controls for LPS, which were stimulated for six hours. However, in typical homeRNA-based studies, collected capillary blood samples would immediately be stabilized. Therefore, the increased inflammatory gene expression response observed over hours would not necessarily translate to typical conditions used in homeRNA studies.

Despite these differences, genome-wide analysis revealed strong concordance between collection methods. The gene expression profiles between unstimulated homeRNA-stabilized samples and unstimulated venous RNA*later*-stabilized showed a high Pearson correlation of 0.948 (Figure S3B), with 15% of detectable genes showing differential expression (Figure S3C). We also performed a similar comparison between stabilization methods (RNA*later* and PAXgene) in the venous blood samples, which showed strong correlation (r=0.948) with 1% of genes demonstrating differential expression (Figure S4), demonstrating that stabilization has minimal impact on the transcriptomic profile of venous blood.

While this gene expression difference in homeRNA-stabilized samples can be partially attributed to differences in collection methodology, we wanted to probe further if we are observing internal variance or between group variance in our samples. We compared technical replicates of RNA*later*-stabilized venous blood from the same donor and these replicates showed a Pearson correlation of 0.933 with no significant DEGs (Figure S5). Since the correlations are similar between the unstimulated homeRNA and Venous RNA*later* samples (r=0.948) and the technical replicates of the Venous RNA*later* samples (r=0.933), there is limited evidence to indicate substantial differences between the capillary (homeRNA) and venous blood samples within the expected error of RNA-sequencing.

These findings demonstrate that while homeRNA-stabilized samples exhibit a baseline inflammatory signature compared to RNA*later*-stabilized venous blood, the two collection methods have highly correlated transcriptomic profiles. Notably, despite this baseline difference, we were still able to detect similar levels of LPS-induced inflammatory response in the stimulated homeRNA samples compared to stimulated venous blood samples (Figure 3), suggesting that the systematic nature of these differences could be accounted for through appropriate normalization strategies in future transcriptomic studies with homeRNA (e.g., normalizing an exposure or targeted response sample with a baseline sample). Further, this baseline inflammatory signature could also be reduced by incorporating an anticoagulant into the Tasso blood collection tube, which has already been incorporated in other Tasso devices such as the Tasso+, which interfaces with any commercially available BD microtainer (including EDTA-coated tubes). Lastly, the high correlation between the unstimulated homeRNA and venous RNA*later* samples observed in this study (r=0.948) is consistent with other comparisons on the gene and protein expression between capillary blood and venous blood^1-5^, supporting the potential of homeRNA to be used in remote transcriptomic studies in place of clinic-based phlebotomy draws.

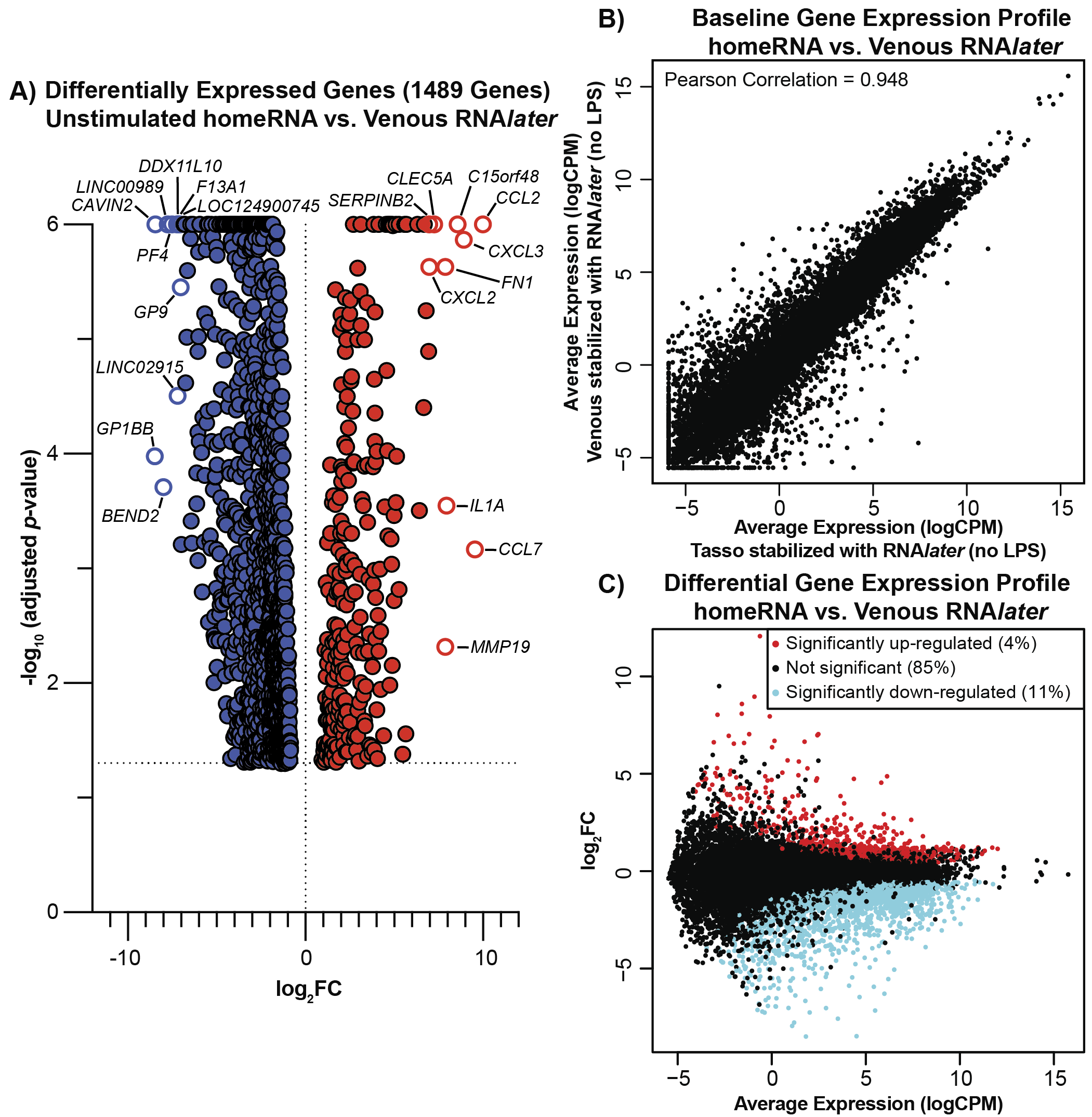

**Figure S3**. Unstimulated homeRNA blood samples exhibit some baseline inflammatory signatures absent in unstimulated RNA*later*-stabilized venous samples while still having highly correlated gene expression profiles. A) Volcano plot of differentially expressed genes (DEGs) between unstimulated homeRNA-stabilized samples and venous blood stabilized with RNA*later*. All DEGs (n=1489 DEGs) are represented by adjusted p-value on the y-axis and log2fold change on the x-axis. Red scatter dots represent up-regulated genes with adjusted p-value ≤ 0.05 and log2fold change ≥ 1.3. Blue scatter dots represent down-regulated with adjusted p-value ≤ 0.05 and log2fold change ≤ -1.3. The top 10 up-regulated and top 10 down-regulated genes are labeled. B) Pearson correlation plot comparing gene expression profiles between homeRNA (Tasso blood stabilized with RNA*later*) and venous RNA*later*-stabilized blood in unstimulated samples. Each point represents a gene, with x-axis showing log counts per million (logCPM) expression in homeRNA samples and y-axis showing logCPM expression in venous RNA*later*-stabilized samples. C) MA-plot showing differential gene expression between homeRNA and venous RNA*later* collection methods. The x-axis represents average expression (logCPM) across samples, and the y-axis shows log2fold change (homeRNA vs. Venous RNA*later*). Red points indicate significantly up-regulated genes (4%) with adjusted p-value ≤ 0.05; blue points indicate significantly down-regulated genes (11%) with adjusted p-value ≤ 0.05; and black points represent non-significant genes (85%).

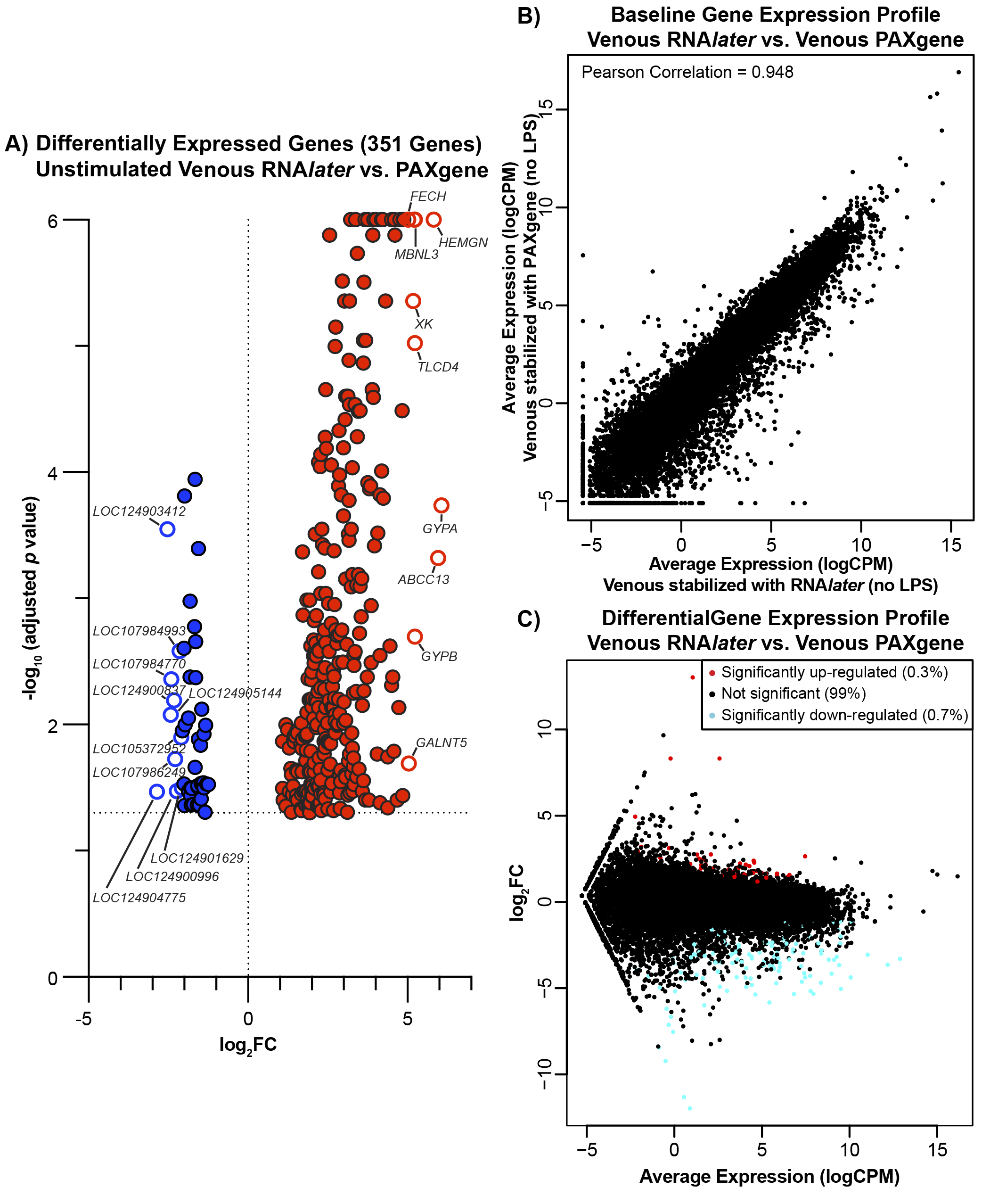

**Figure S4**. Comparison of gene expression signature in unstimulated (no LPS) venous blood stabilized with RNA*later* and PAXgene. A) Volcano plot of differentially expressed genes (DEGs) between unstimulated venous RNA*later*-stabilized samples and venous PAXgene-stabilized samples. All DEGs (n=351 DEGs) are represented by adjusted p-value on the y-axis and fold change on the x-axis. Red scatter dots represent up-regulated genes with adjusted p-value ≤ 0.05 and log2fold change ≥ 1.3. Blue scatter dots represent down with adjusted p-value ≤ 0.05 and log2fold change ≤ -1.3. The top 10 up-regulated and top 10 down-regulated genes are labeled. B) Pearson correlation plot comparing gene expression profiles between venous blood stimulated with RNA*later* and PAXgene in unstimulated samples. Each point represents a gene, with x-axis showing log counts per million (logCPM) expression in venous RNA*later*-stabilized samples and y-axis showing logCPM expression in venous PAXgene-stabilized samples. C) MA-plot showing differential gene expression between RNA*later* and PAXgene stabilization methods. The x-axis represents average logCPM across samples, and the y-axis shows log2fold change (venous RNA*later* vs. venous PAXgene). Red points indicate significantly upregulated genes (0.3%) with adjusted p-value ≤ 0.05; blue points indicate significantly downregulated genes (0.7%) with adjusted p-value ≤ 0.05; black points represent non-significant genes (99%).

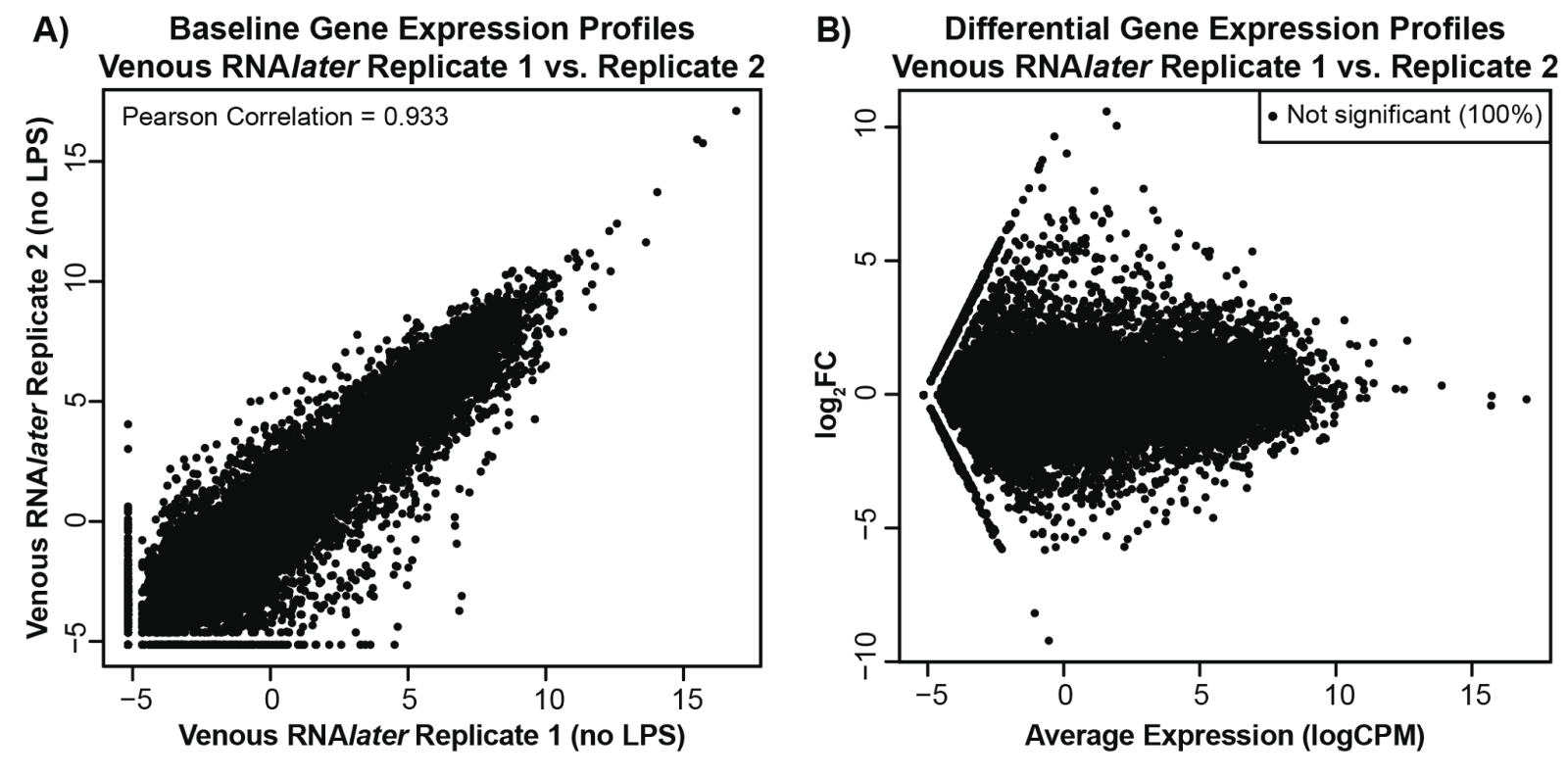

**Figure S5**: Technical variability in venous RNA*later*-stabilized blood samples. A) Pearson correlation plot comparing gene expression profiles between two unstimulated technical replicates of venous RNA*later*-stabilized samples from all three donors. Each point represents a gene, with x-axis showing log counts per million (logCPM) expression in replicate 1 and y-axis showing logCPM expression in replicate 2. C) MA-plot showing differential gene expression between replicates 1 and 2 in venous RNA*later*-stabilized blood samples from all three donors. The x-axis represents average expression (logCPM) across samples, and the y-axis shows log_2_fold change (replicate 1 vs. replicate 2). Black points represent non-significant genes (100%).

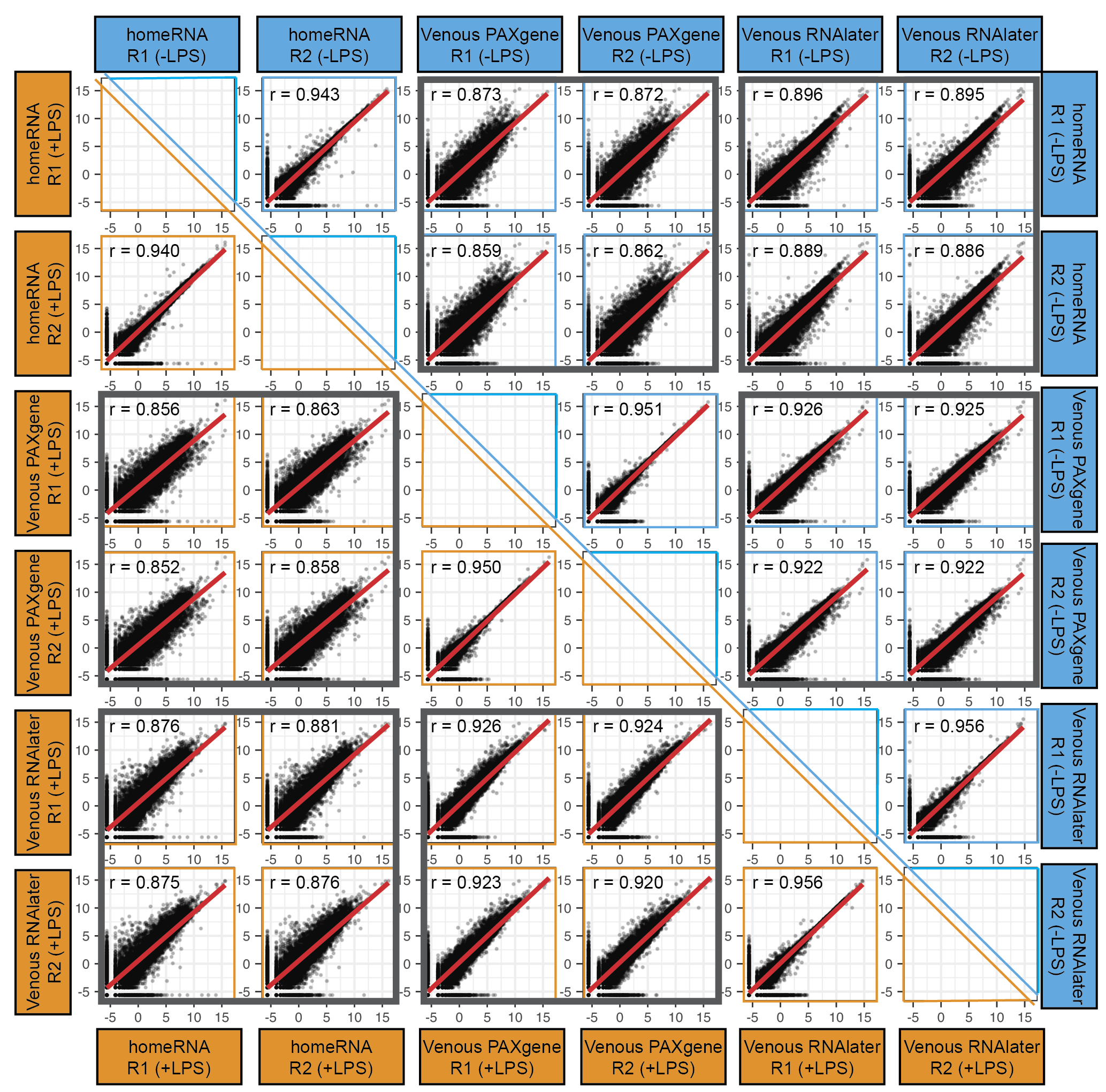

**Figure S6.** LPS-stimulated (orange) and unstimulated (blue) gene expression profiles are similar between homeRNA-stabilized capillary blood, RNA*later*-stabilized venous blood, and PAXgene-stabilized blood samples in Donor 2. Scatter plots compare log2-transformed normalized gene expression values (logCPM) between replicates of homeRNA capillary blood (Tasso-collected and RNA*later*-stabilized), PAXgene-stabilized venous blood, and RNA*later*-stabilized venous blood samples. Unstimulated samples (-LPS) are plotted in the upper right triangle (blue panels) and LPS-stimulated samples (+LPS) are plotted in the lower left triangle (orange panels). Each sample was collected and processed in technical replicates (R1 and R2). Within-method comparisons show high technical reproducibility between replicates for homeRNA (r = 0.943 for -LPS; r = 0.940 for +LPS), Venous PAXgene (r = 0.951 for -LPS; r = 0.950 for +LPS), and Venous RNA*later* (r = 0.956 for -LPS; r = 0.956 for +LPS). Between-method comparisons (boxed in gray) show strong correlations across all collection and stabilization approaches under both unstimulated conditions (homeRNA vs. Venous PAXgene: r = 0.859 to 0.873; homeRNA vs. Venous RNA*later*: r = 0.886 to 0.896; Venous PAXgene vs. Venous RNA*later*: r = 0.922 to 0.926) and LPS-stimulated conditions (homeRNA vs. Venous PAXgene: r = 0.852 to 0.863; homeRNA vs. Venous RNA*later*: r = 0.875 to 0.886; Venous PAXgene vs. Venous RNA*later*: r = 0.920 to 0.926). Red lines indicate the line of best fit; Pearson correlation coefficients (r) are displayed in each panel.

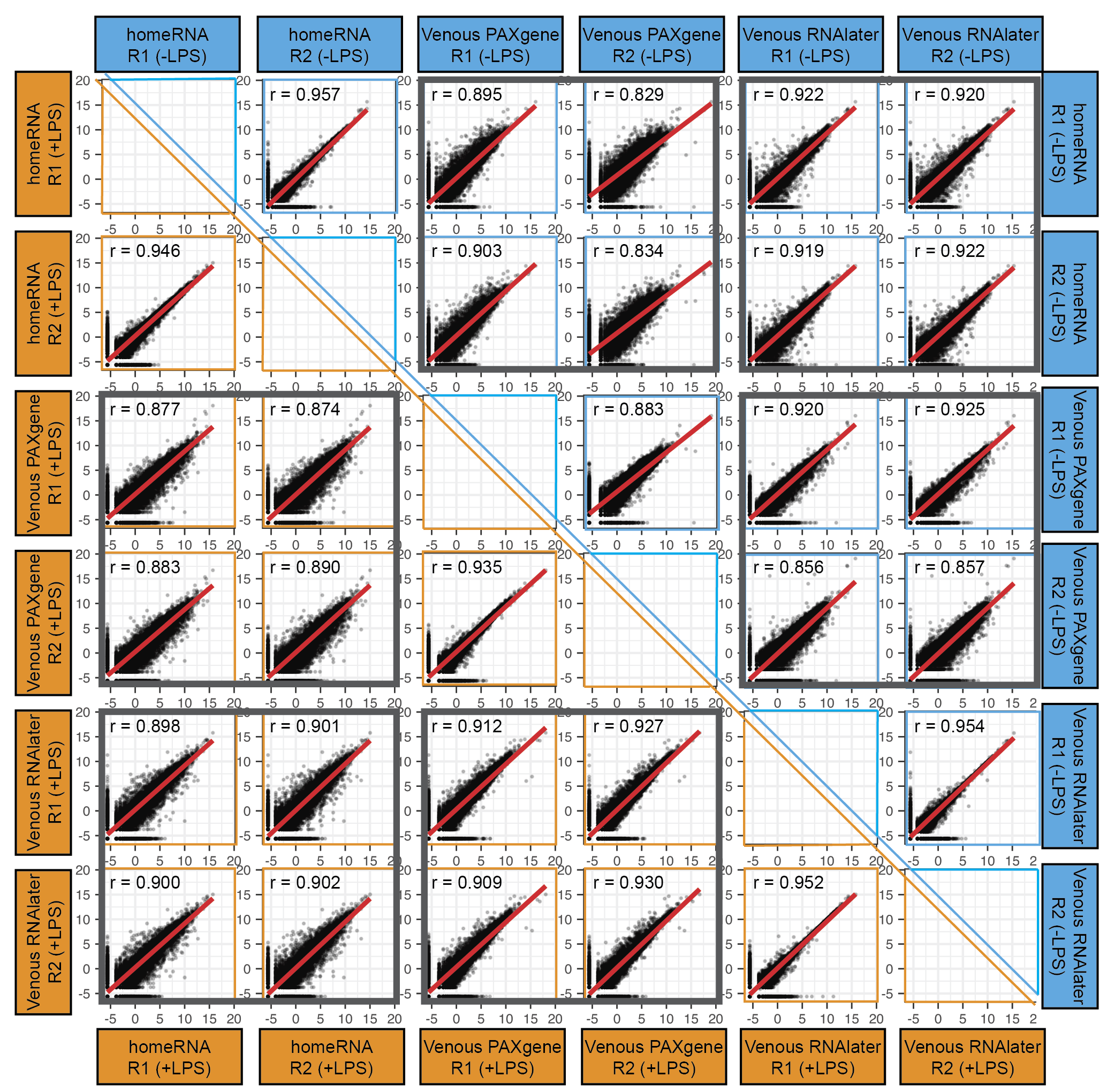

**Figure S7.** LPS-stimulated (orange) and unstimulated (blue) gene expression profiles are similar between homeRNA-stabilized capillary blood, RNA*later*-stabilized venous blood, and PAXgene-stabilized blood samples in Donor 3. Scatter plots compare log2-transformed normalized gene expression values (logCPM) between replicates of homeRNA capillary blood (Tasso-collected and RNA*later*-stabilized), PAXgene-stabilized venous blood, and RNA*later*-stabilized venous blood samples. Unstimulated samples (-LPS) are plotted in the upper right triangle (blue panels) and LPS-stimulated samples (+LPS) are plotted in the lower left triangle (orange panels). Each sample was collected and processed in technical replicates (R1 and R2). Within-method comparisons show high technical reproducibility between replicates for homeRNA (r = 0.957 for -LPS; r = 0.946 for +LPS), Venous PAXgene (r = 0.883 for -LPS; r = 0.935 for +LPS), and Venous RNA*later* (r = 0.954 for -LPS; r = 0.952 for +LPS). Between-method comparisons (boxed in gray) show strong correlations across all collection and stabilization approaches under both unstimulated conditions (homeRNA vs. Venous PAXgene: r = 0.829 to 0.903; homeRNA vs. Venous RNA*later*: r = 0.920 to 0.922; Venous PAXgene vs. Venous RNA*later*: r = 0.856 to 0.925) and LPS-stimulated conditions (homeRNA vs. Venous PAXgene: r = 0.874 to 0.890; homeRNA vs. Venous RNA*later*: r = 0.898 to 0.902; Venous PAXgene vs. Venous RNA*later*: r = 0.909 to 0.930). Red lines indicate the line of best fit; Pearson correlation coefficients (r) are displayed in each panel.
